# Molecular layer interneurons shape the spike activity of cerebellar Purkinje cells

**DOI:** 10.1101/376517

**Authors:** Amanda M. Brown, Marife Arancillo, Tao Lin, Daniel R. Catt, Joy Zhou, Elizabeth P. Lackey, Trace L. Stay, Zhongyuan Zuo, Joshua J. White, Roy V. Sillitoe

## Abstract

**One-sentence summary:** Cerebellar stellate cells and basket cells shape distinct Purkinje cell firing properties

**Abstract:** Purkinje cells receive synaptic input from several classes of interneurons. Here, we address the roles of inhibitory molecular layer interneurons in establishing Purkinje cell function *in vivo.* Using conditional genetics approaches in mice, we compare how the lack of stellate cell versus basket cell GABAergic neurotransmission sculpts the firing properties of Purkinje cells. We take advantage of an inducible *Ascl1^CreER^* allele to spatially and temporally target the deletion of the vesicular GABA transporter, *Vgat*, in developing neurons. Selective depletion of basket cell GABAergic neurotransmission increases the frequency of Purkinje cell simple spike firing and decreases the frequency of complex spike firing in adult behaving mice. In contrast, lack of stellate cell communication increases the regularity of Purkinje cell simple spike firing while increasing the frequency of complex spike firing. Our data uncover complementary roles for molecular layer interneurons in shaping the rate and pattern of Purkinje cell activity *in vivo.*

## Introduction

The cerebellum is essential for diverse motor functions including coordination, learning, posture, and balance (Manto et al., 2012). Despite this functional diversity, a core cerebellar circuit mediates all of its functions (Eccles, 1967; Reeber et al., 2013). This canonical cerebellar circuit is comprised of relatively few types of cells (Voogd and Glickstein, 1998). The Purkinje cells, the sole output of the cerebellar cortex and main computational cell type, are located at the center of the circuit (Figure 1A). Purkinje cells receive input from several classes of interneurons. The granule cells project parallel fibers that send excitatory signals to Purkinje cells (Barbour, 1993; Eccles et al., 1966a, 1966b; Konnerth et al., 1990). However, in the posterior cerebellum, the unipolar brush cell interneurons can influence granule cell output by amplifying vestibular inputs that are delivered to the cerebellum by mossy fibers (Mugnaini et al., 2011). Golgi cells, another cell type of the granular layer, provide feedforward and feedback inhibitory signals onto granule cells (Cesana et al., 2013; Hull and Regehr, 2012). Purkinje cells also receive direct inhibitory inputs from basket cells that form pericellular baskets as well as specialized terminals known as pinceaux, and also from stellate cells that terminate on the smooth shafts of the Purkinje cell dendrites (Eccles et al., 1965; Figure 1B-C). Together, the interneurons play an essential role in controlling cerebellar cortical output during motor behavior (Barmack and Yakhnitsa, 2008). However, how each class of interneurons influences Purkinje cell firing is poorly understood. Here, we used inducible conditional genetic approaches in mice to test whether the two classes of cerebellar molecular layer interneurons have dedicated GABAergic functions *in vivo.*

**Figure 1.**
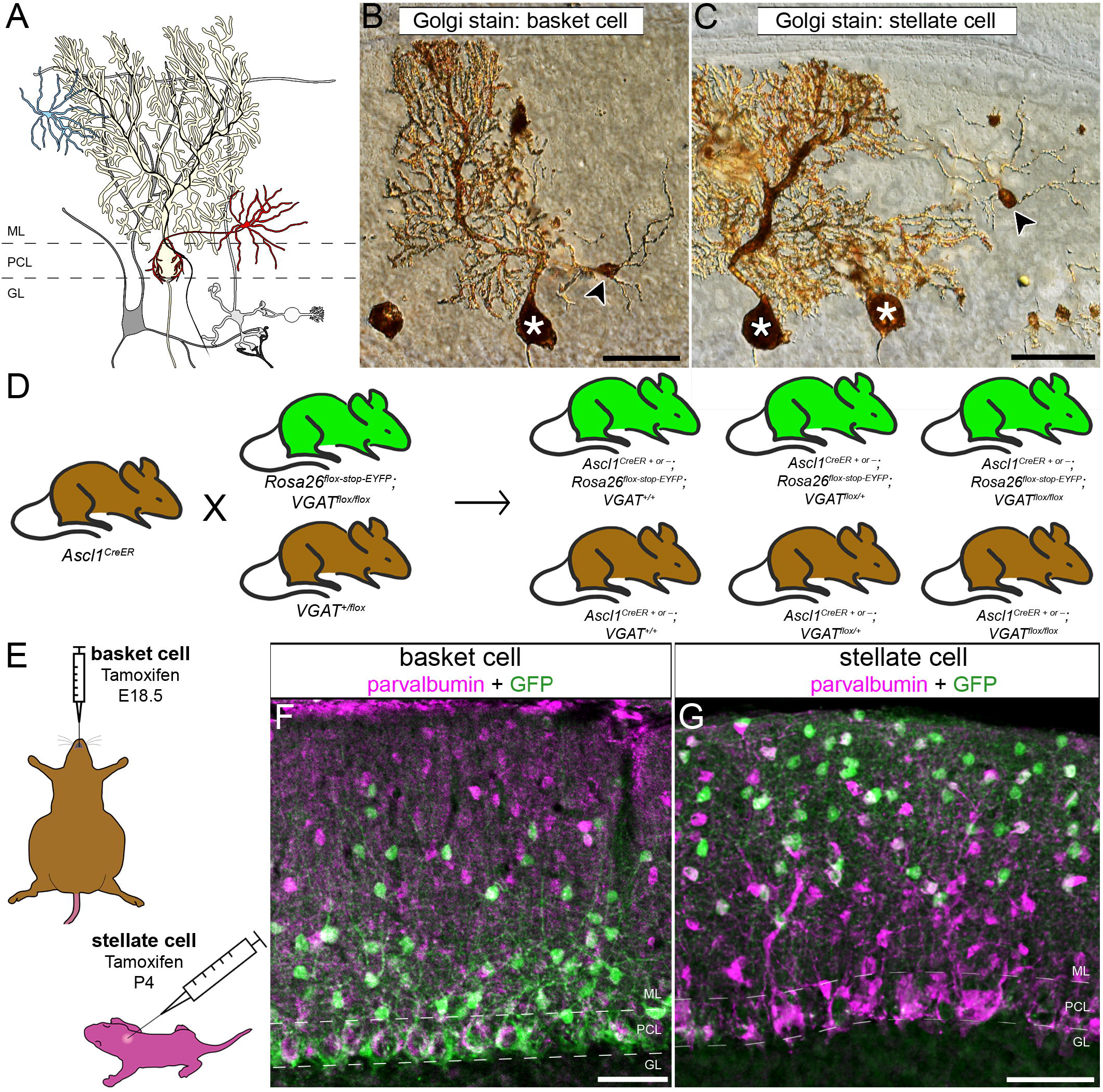
The *Ascl1^CreER^* allele can be used for genetic marking of stellate cells and basket cells. (**A**) Schematic of cerebellar circuitry. Purkinje cell (yellow), basket cell (red), and stellate cell (blue) are colorized while other cells and fibers in the cerebellar cortex are represented in grayscale. Dotted lines represent the borders of the Purkinje cell layer (PCL) with the molecular layer (ML) and granule cell layer (GL). (**B-C**) Golgi-Cox stain of cerebellar tissue. (**B**) Basket cell (arrowhead) and Purkinje cell (asterisk) revealed by Golgi-Cox stain. Scale = 50μm. (**C**) Stellate cell (arrowhead) and Purkinje cells (asterisks) revealed by Golgi-Cox stain. Scale = 50μm. (**D**) Representation of breeding scheme. (**E**) Schematic of methods for tamoxifen administration. Tamoxifen was administered via oral gavage to pregnant dams at E18.5 to achieve constitutive marking and manipulation of a subset of basket cells in the resulting pups (upper left). Tamoxifen was administered via subcutaneous injection into the scruff of pups at P4 to achieve constitutive marking and manipulation of a subset of stellate cells (bottom right). (**F-G**) Sagittal cerebellar sections from tamoxifen-treated animals stained with parvalbumin, a marker of inhibitory cerebellar neurons including Purkinje cells and molecular layer interneurons, and GFP to highlight the genetically marked cells. Scale = 50μm. (**F**) Sagittal cerebellar section from an animal treated with tamoxifen at the basket cell marking time point. (**G**) Sagittal cerebellar section from an animal treated with tamoxifen at the stellate cell marking time point.

Cerebellar interneurons come from distinct lineages and have specific birth dates (Machold and Fishell, 2005; Maricich and Herrup, 1999; Wang et al., 2005; Zhang and Goldman, 1996). Fate mapping and transplant experiments demonstrated that the inhibitory interneurons are generated in a precise spatial and temporal manner such that the early born neurons occupy deep positions within the cerebellar cortex whereas later born neurons migrate to the more superficial locations (Altman and Bayer, 1997; Leto et al., 2009; Weisheit et al., 2006). More recent genetic inducible fate mapping experiments corroborated those results, and further suggested that the timing of *Ascl1* gene expression during differentiation may be used as a molecular time stamp for the birth of specific classes of GABAergic interneurons (Sudarov et al., 2011). *Ascl1*, also known as *Mash1*, is a basic helix-loop-helix transcription factor that is expressed during cerebellar development (Kim et al., 2008; Sudarov et al., 2011). In this study, we used the *Ascl1^CreER^* genetic fate-mapping allele (Sudarov et al., 2011) to not only mark interneurons, but also to constitutively silence their output. To do so, we selectively deleted a critical functional domain in the *Vgat* gene (Tong et al., 2008), which removed the ability of the inhibitory interneurons to signal their output using fast GABAergic neurotransmission. Genetic deletion using *Ascl1^CreER^* allowed us to independently target newly differentiated stellate and basket cell interneurons in the molecular layer because these neurons are born almost exclusively during the peri-to postnatal period when the cerebellar circuits are wiring up for function (White and Sillitoe, 2013b). This is advantageous for our study because *in vitro* studies showed that as development progresses, interneuron to Purkinje cell inhibition increases (Pouzat and Hestrin, 1997). Functional studies support these data since removing the interneurons or their postsynaptic γ2 GABA(A) receptors obstruct motor learning (Sergaki et al., 2017; Wulff et al., 2009). Recent work also demonstrates that movement rate is dependent on coordinated molecular layer interneuron activity (Gaffield and Christie, 2017). Still, there is a long-standing debate as to whether stellate cells and basket cells are distinct types of interneurons (Schilling and Oberdick, 2009; Sultan and Bower, 1998), and more broadly whether they perform different functions in the cerebellar circuit (He et al., 2015). In this study, we genetically mark stellate cells and basket cells independently and manipulate their GABAergic neurotransmission as the cells are born to determine their impact on establishing the mature firing properties of Purkinje cells *in vivo.*

## Results

### A mouse genetic strategy for marking and manipulating cerebellar GABA interneurons

We aimed to manipulate neurotransmission in a way that would block the activity of the molecular layer interneurons without inducing changes in cerebellar morphology or causing neurodegeneration. We therefore targeted the function of the vesicular GABA transporter (VGAT), a transporter that is essential for the uptake of GABA into synaptic vesicles. Conditional knockout of *Vgat* in neurons does not induce widespread defects in cerebellar anatomy (White et al., 2014), making it an ideal target for genetic deletion. We targeted the removal of the *Vgat* gene in stellate cells and basket cells in the cerebellar cortex by using the *Mash1/Ascl1* promoter to drive tamoxifen-inducible Cre in the cerebellum (Sudarov et al., 2011; Figure 1D). The *Mash1/Ascl1* gene (referred to from here on as *Ascl1)* encodes a developmental transcription factor that is critical for the specification of neurons and glia (Kim et al., 2008). In the cerebellum, it is expressed in waves by neural and glial precursors as cells exit the cell cycle and begin to differentiate (Sudarov et al., 2011). The period of stellate cell differentiation begins at late embryonic stages and reaches peak levels at postnatal day (P) 3 - P5 whereas basket cell differentiation occurs during late embryogenesis and peaks at around embryonic day (E) 18 (Sudarov et al., 2011). Therefore, to specifically target stellate cells we subcutaneously injected *Ascl1^CreER^;R26^fx-stop-EYFP^* postnatal pups with a single 20 mg/ml dose of tamoxifen at P4 (Figure 1E and G), which would allow for recombination in *Ascl1* expressing cells for the next ~32 hours (Zervas et al., 2004). But note that we predicted to label only subsets of interneurons since they are born over several days. Analysis of the GFP expression showed labeling of neurons in the upper two thirds of the molecular layer (Figure 1G). Morphological analysis of individual neurons that were marked by GFP confirmed their “stellate” appearance as well as their pattern of axonal projections within the molecular layer (Figure 1G and 2A). We next confirmed whether we could target putative basket cells, as demonstrated previously using a different reporter (Sudarov et al., 2011). We targeted the reporter to neurons located in the basal one third of the molecular layer by delivering tamoxifen to E18.5 embryos by oral gavage of *Ascl1^CreER^;R26^fx-stop-EYFP^* pregnant dams (Figure 1E and F). The morphology of these neurons was consistent with their identity as basket cells, namely because of the presence of baskets on the Purkinje cell somata (Figure 1F and 2A). We could also track their prominent axons that travel in a transverse trajectory within the molecular layer, in close proximity to their targets, the Purkinje cell somata, which are located immediately below the axons (Figure 1F and 2A).

**Figure 2.**
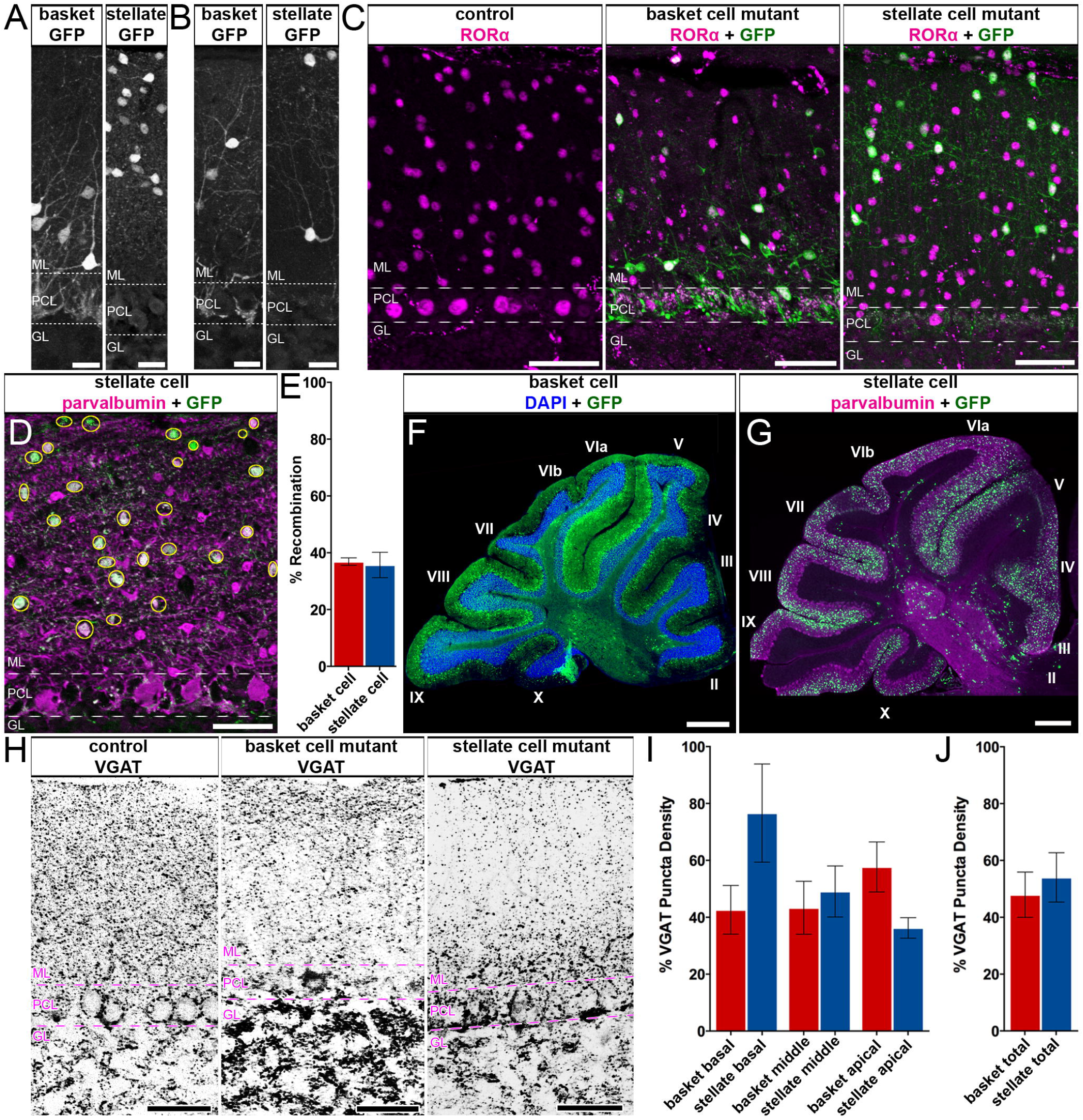
*Ascl1^CreER^* conditional deletion of VGAT protein is efficient and selective. (**A-B**) Sagittal cerebellar sections stained with GFP to reveal labeled basket and stellate cells. (**A**) Labeled basket cell somas are predominantly found in the basal molecular layer (ML) and their processes form conspicuous baskets around Purkinje cell somas in the Purkinje cell layer (PCL) (left). Labeled stellate cell somas are principally found in the apical ML with no processes forming baskets around Purkinje cell somas (right). Scale = 20μm. (**B**) Some labeled basket cells were found in more apical regions of the ML, however their processes still descended through the ML to form baskets around Purkinje cell somas (left). Some labeled stellate cells were found towards basal regions of the ML, however their processes ascended to the apical ML and did not form baskets around Purkinje cell somas (right). Scale = 20μm. (**C**) Sagittal cerebellar tissue with ML interneuron somas stained with RORα and genetically labeled cells stained with GFP. Control tissue stained with RORα shows RORα expression in some Purkinje cells and uniformly throughout the ML in molecular layer interneurons (left). RORα expression is unaltered in basket cell (middle) and stellate cell (right) silenced mutant mice. Scale = 50μm. (**D**) Representation of recombination quantification in a sagittal cerebellar section with labeled stellate cells. Inhibitory interneurons including Purkinje cells and ML interneurons are stained with parvalbumin. Labeled cells are stained with GFP and counted (yellow circles). Scale = 50μm. (**E**) Quantification of recombination efficiency in basket and stellate cell conditions (basket cell recombination: 34.26% ± 1.316%; stellate cell recombination: 35.69% ± 4.458%). (**F-G**) Sample of a whole sagittal cerebellar section in the basket (**F**) and stellate (**G**) manipulation conditions. Scale = 0.5mm. (**F**) In the basket cell marking condition, granule cells are highlighted with heavy DAPI staining and marked cells are found throughout the section with no obvious patterning and close to the granule cell layer in the basal ML. (**G**) In the stellate cell marking condition, Purkinje cells and ML interneurons are stained with parvalbumin. Labeled cells are stained with GFP and found in the apical molecular layer throughout the section with no obvious patterning. (**F-G**) Cerebellar lobules are indicated with Roman numerals. Scale = 0.5mm. (**H**) Sagittal cerebellar tissue stained with VGAT to mark inhibitory synapses. VGAT expression was uniform across the ML in control mice (left), but was significantly reduced in the basal and middle ML in basket cell VGAT deletion mice (middle) and significantly reduced in the apical ML in stellate cell VGAT deletion mice (right). Scale = 50μm. (**I**) Quantification of VGAT puncta density in the basal, middle, and apical ML of basket and stellate VGAT mutant mice (basket cell mean VGAT density as percent of control: apical = 57.69% ± 8.799, middle = 43.33% ± 9.317, basal = 42.62% ± 8.560; stellate cell mean VGAT density as percent of control: apical = 36.26% ± 3.621, middle = 49.08% ± 8.957, basal = 76.63% ± 17.23). (**J**) Quantification of VGAT puncta density throughout the entire ML of basket and stellate VGAT mutant mice. There is no significant difference in the density of VGAT puncta between the two mutant conditions (basket cell mean VGAT density as percent of control = 47.95 ± 7.962; stellate mean VGAT density as percent of control = 54.03 ± 8.703; *P* = 0.6336). (**A-D, H**) Dotted lines indicate the borders of the Purkinje cell layer (PCL) with the molecular layer (ML) above and the granule layer (GL) below.

In addition to labeling what would be considered typical stellate cells and basket cells (Figure 2A), we could also reveal neurons with structural variations, but likely belonging to these same classes. In the stellate cell marking scheme, cells with a more restricted dendritic span were observed in the very apical regions of the molecular layer (Figure 2A), and within the middle of the layer we could label cells with soma positions that mimicked basket cells (Figure 2B). Regardless of soma position, in the stellate cell marking scheme, the predominant loss of VGAT expression in the deletion allele was always in the more apical locations of the molecular layer (see Figure 2H). The basket cell marking scheme also labeled cells in the middle portion of the molecular layer, and these cells projected either ascending or descending processes (Figure 2A). Therefore, although the stellate cells and basket cells, defined strictly on position and density, could be separated using the *Ascl1^CreER^* lineage tracing, each class also contains cells with a varying range of specializations that are observed in their dendritic processes and axonal projections. Thus, neuronal position within the molecular layer alone is not necessarily indicative of the identity of that interneuron, or the specific interneuron cell class that it belongs to. However, the cellular anatomy revealed by our genetic marking data are consistent with the results of classic Golgi staining of molecular layer interneurons (Palay and Chan-Palay, 1974).

### The *Ascl1^CreER^* allele has high specificity and recombination efficiency in interneurons

We next tested whether we could confirm if the labeling of apical and basal molecular layer neurons reflect specifically stellate cell and basket cells, respectively. The reporter expressing cells colocalized with the expression of parvalbumin, which is a well-known marker for Purkinje cells and molecular layer interneurons (Stichel et al., 1986; Figure 1F, 1G, and 2D). We did not detect any GFP labeling in parvalbumin-immunoreactive Purkinje cells (Figure 1F, 1G, and 2D), which is consistent with the earlier marking of Purkinje cells with *Ascl1^CreER^* between E10 and E13 (Sudarov et al., 2011). The distribution of reporter expression in stellate versus basket cells was validated by RAR-related orphan receptor alpha (RORα) expression (Figure 2C), which also marks molecular layer interneurons and Purkinje cells (Hamilton et al., 1996; Ino, 2004; Maricich and Herrup, 1999; Sillitoe et al., 2009). An advantage of using RORα expression, a nuclear hormone receptor, is that the cytoplasmic GFP labeling in marked neurons pairs nicely with the robust staining of the nucleus. The adult stellate cells that were marked by giving pups tamoxifen at P4 expressed RORα, as did the adult basket cells that were marked at E18.5 (Figure 2C middle and right). Similar to parvalbumin, when we used RORα expression as a marker we did not detect GFP in Purkinje cells (Figure 2C **middle and right**). Moreover, we did not detect GFP expression in any of the granular layer interneurons (Figure 2C **middle and right**). We conclude that our *Ascl1^CreER^* genetic marking schemes are selective for the classes of inhibitory interneurons that reside within the molecular layer. With consideration of these classes’ date of differentiation, morphology, layer location, and protein expression profile, for the remaining duration of this text, the cells that are marked using the E18.5 and P4 induction time points will be referred to as “basket cells” and “stellate cells,” respectively.

The efficiency of *Ascl1^CreER^* recombination on the *R26^fx-stop-EYFP^* reporter allele provides a prediction for the percentage of interneurons that can be manipulated with this genetic paradigm. It was essential to know how widespread and reliable the cell marking strategy is before crossing the *Ascl1^CreER^* line to a functional allele such as *Vgat^fx/fx^* for testing circuit function. We quantitatively examined the number of parvalbumin-expressing molecular layer interneurons that also express GFP reporter in *Ascl1^CreER^;R26^fx-stop-EYFP^* mice (Figure 2D). The recombination observed in the totality of the molecular layer in the stellate cell marking scheme is 35.69% ± 4.458% with the vast majority of labeled cells observed in the apical molecular layer (Figure 2E, see 1G **and** 2C **right**; n=2 sections from 3 animals each). Similarly, the recombination observed in the totality of the molecular layer in the basket cell scheme is 34.26% ± 1.316% with the majority of labeled cells located in the basal and middle molecular layer (Figure 2E, **see** 1F and 2C **middle**; n=3 sections from 3 animals each). This percent recombination marked enough putative basket cells to project axons that form baskets on almost every Purkinje cell in the field of view, on any given tissue section (Figure 1F, 2C **middle**). Importantly, we were successful in manipulating a similar number of cells in both conditions while avoiding unwanted recombination throughout the entire molecular layer, which was required for targeting each class of interneurons. This marking is ideal for distinguishing the relative distributions of neuron that contribute to the molecular layer populations. Additionally, in both genetic marking paradigms, we could detect GFP reporter expression in all lobules of the cerebellum, and we were able to mark neurons in the vermis, paravermis, and also in the hemispheres (Figure 2F-G). Therefore, we did not find systematic regional biases in the localization of interneuron populations that were targeted by our genetic marking paradigms.

### Targeted loss of VGAT protein in conditional *Ascl1^CreER/+^; Vgat^fx/fx^* mutant mice

We set up crosses to generate litters with genotype *Ascl1^CreER/+^;R26^fx-stop-EYFP^; Vgat^+/+^* (control) and genotype *Ascl1^CreER/+^; R26^fx-stop-EYFP^;Vgat^fx/fx^* (mutant). This approach allows us to mark and manipulate the same neurons *in vivo.* After tamoxifen treatment at P4, we expected to mark and manipulate stellate cells in the mutants and only mark cells in the control. We expected a similar manipulation for basket cells after providing tamoxifen at E18.5. To test whether VGAT was removed from the intended neurons we quantified the number and distribution of VGAT-positive synaptic terminals in the molecular layer. VGAT expression in the molecular layer of control mice showed an approximately uniform distribution of punctae from the basal to the apical regions (Figure 2H). Stellate cells, basket cells, and Purkinje cell axon collaterals are the main contributors to the GABAergic synapses marked by VGAT expression in the molecular layer. We found that the density of VGAT punctae in the stellate cell mutant mice was significantly reduced, specifically in the apical region of the molecular layer (Figure 2H and 2I; mean VGAT density as percent of control: apical = 36.26% ± 3.621, *P* = 0.0249; middle = 49.08% ± 8.957, *P* = 0.0740; basal = 76.63% ± 17.23, *P* = 0.4320). In contrast, after deleting *Vgat* in basket cells, we found significantly reduced expression of VGAT in the basal portion of the molecular layer, but we also found a marked reduction, albeit less pronounced, in the middle and apical regions (Figure 2H and 2I; mean VGAT density as percent of control: apical = 57.69 *%* ± 8.799, *P* = 0.0976; middle = 43.33% ± 9.317, *P* = 0.0451; basal = 42.62% ± 8.560, *P* = 0.0315). Loss of basal VGAT expression is due to manipulation of the baskets and pinceaux whereas loss of VGAT apically is due to manipulation of basket cell synapses made by the ascending collateral axons (Palay and Chan-Palay, 1974; Figure 2H, **see** 2A **and** 1B). Interestingly, total VGAT expression in the molecular layer was not significantly different between the basket cell and stellate cell manipulations (Figure 2J; basket cell mean VGAT density as percent of control = 47.95 ± 7.962; stellate mean VGAT density as percent of control = 54.03 ± 8.703; *P* = 0.6336). These data confirm that genetic deletion of *Vgat* with *Ascl1^CreER^* is effective for manipulating VGAT protein. The data also show that the *Ascl1^CreER^* allele can be used for region-specific deletion of VGAT in a cerebellar layer where classes of related neurons are co-residing.

### Deletion of *Vgat* does not prevent interneurons from occupying the molecular layer

Deletion of *Vgat* could result in a loss of VGAT because the protein is depleted or because cells are lost. Indeed deletion of genes encoding for molecules involved in neurotransmission can result in cerebellar cell death, especially when these molecules are expressed during development (McFarland et al., 2007; Sawada et al., 2009; Slemmer et al., 2005). To test this possibility, we again stained for the nuclear hormone receptor, RORα, to visualize interneuron distribution in lobule III or IV. Lobules III and IV are ideal for systematically examining molecular layer anatomy because the deep fissures provide long, straight regions of cortex that allow consistent measures for analysis. We found that the density of molecular layer interneurons that express RORα in both the stellate cell and basket cell mutants (Figure 2C; stellate cells – control = 1.215×10^−4^ cells/μm^3^ ± 3.604×10^−5^, N = 3, n = 3; mutant = 1.168×10^−4^ cells/μm^3^ ± 1.711 x10^−5^; *P* = 0.9135, N = 3, n = 3; basket cells – control = 1.141×10^−4^ cells/μm^3^ ± 1.137 x10^−5^, N = 3, n = 3; mutant = 9.559×10^−5^ cells/μm^3^ ± 2.210×10^−5^, N = 3, n = 3; *P* = 0.5098) was not significantly different from controls. Therefore, loss of VGAT does not kill the interneurons.

### Loss of *Vgat* in newly differentiated interneurons causes Purkinje cell firing defects

To test for electrophysiology defects we analyzed *Ascl1^CreER/+^;R26^fx-stop-EYFP^;Vgat^+/+^* (control) and *Ascl1^CreER/+^; R26^fx-stop-EYFP^;Vgat^fx/fx^* mice (mutant). However, *Ascl1^CreER/+^;Vgat^fx/fx^* mutants without the marking allele were also used for analysis. We performed extracellular single-unit recordings with tungsten electrodes. To access the cerebellum, a craniotomy and recording port were positioned over lobule VI of the vermis (Figure 3A; White et al., 2016a). Alert adult mice were allowed to stand on a wheel during recordings (Figure 3B). Although the mice are free to walk on the wheel, the periods of most stable recordings that were used to quantify the Purkinje cell responses were acquired when the mice were sitting at rest. Purkinje cells were recorded at a depth of 0 – 2 mm from the surface of the cerebellum and were identified by their characteristic complex spikes (Figure 3C). To examine the firing properties of Purkinje cells, we measured both the firing frequency and the variability of the firing pattern in alert mice for both simple spike and complex spike activity. Firing frequency was measured as the mean number of spikes over time, and indicates the level of activity of a cell. The variability of the firing pattern was measured using two parameters: the coefficient of variance (CV), which measures the variability in firing intervals over the entire recording session, and CV2, which measures the variability of firing intervals between two adjacent spikes (Holt et al., 1996). Loss of stellate cell GABAergic neurotransmission increases the regularity of Purkinje cell simple spike firing as measured by CV2 (Figure 3G; control = 0.5186 ± 0.01968; N = 7, n = 20; mutant = 0.4091 ± 0.01220; N = 3, n = 15; *P* < 0.0001). However, we did not detect a significant change in CV (Figure 3F; control = 0.5787 ± 0.0264; N = 7, n = 20; mutant = 0.5333 ± 0.03603; N = 3, n = 15; *P* = 0.3181) or the firing rate (Figure 3E; control = 74.99Hz ± 6.031Hz; N = 7, n = 20; mutant = 76.85Hz ± 8.298Hz; N = 3, n = 15; *P* = 0.8577). Interestingly, loss of basket cell GABAergic neurotransmission resulted in an increase in the frequency of Purkinje cell simple spike firing (Figure 3M; control = 64.56Hz ± 4.615Hz; N = 5, n = 17; mutant = 84.76Hz ± 5.670Hz; N = 3, n = 18; *P* = 0.0094). There was no significant change in CV (Figure 3N; control = 0.5996 ± 0.03410; N = 5, n = 17; mutant = 0.5611 ± 0.03852; N = 3, n = 18; *P* = 0.4599) or CV2 (Figure 3O; control = 0.5223 ± 0.02673; N = 5, n =17; mutant = 0.4688 ± 0.03437; *P* = 0.2282). Further, there was a divergent effect of the lack of stellate and basket cell GABAergic neurotransmission on complex spike activity. Lack of stellate cell neurotransmission increases the complex spike firing rate (Figure 3I; control = 1.186Hz ± 0.06845Hz; N = 7, n = 20; mutant = 1.469Hz ± 0.08279Hz; N = 3, n = 15; *P* = 0.0131). This occurs without a significant change in CV (Figure 3J; control = 0.8412 ± 0.03723; N = 7, n = 20; mutant = 0.7650 ± 0.02704; N = 3, n = 15; *P* = 0.1076) or CV2 (Figure 3K; control = 0.8819 ± 0.02203; N = 7, n = 20; mutant = 0.8456 ± 0.01697; N = 3, n = 15; *P* = 0.2006). However, the lack of basket cell neurotransmission decreases the complex spike firing rate (Figure 3Q; control = 1.538Hz ± 0.07298Hz; N = 5, n = 17; mutant = 1.148Hz ± 0.03695Hz; N = 3, n = 18; *P* < 0.0001). This also occurs without a significant change in CV (Figure 3R; control = 0.7074 ± 0.01833; N = 5, n = 17; mutant = 0.7342 ± 0.01970; N = 3, n = 18; *P* = 0.3252) or CV2 (Figure 3S; control = 0.8533 ± 0.01476; N = 5, n = 17; mutant = 0.8552 ± 0.02171; N = 3, n = 18; *P* = 0.9414). These data suggest that stellate cell and basket cell GABAergic output activity cooperate to establish the proper rate and pattern of simple spike and complex spike firing of Purkinje cells *in vivo.*

**Figure 3.**
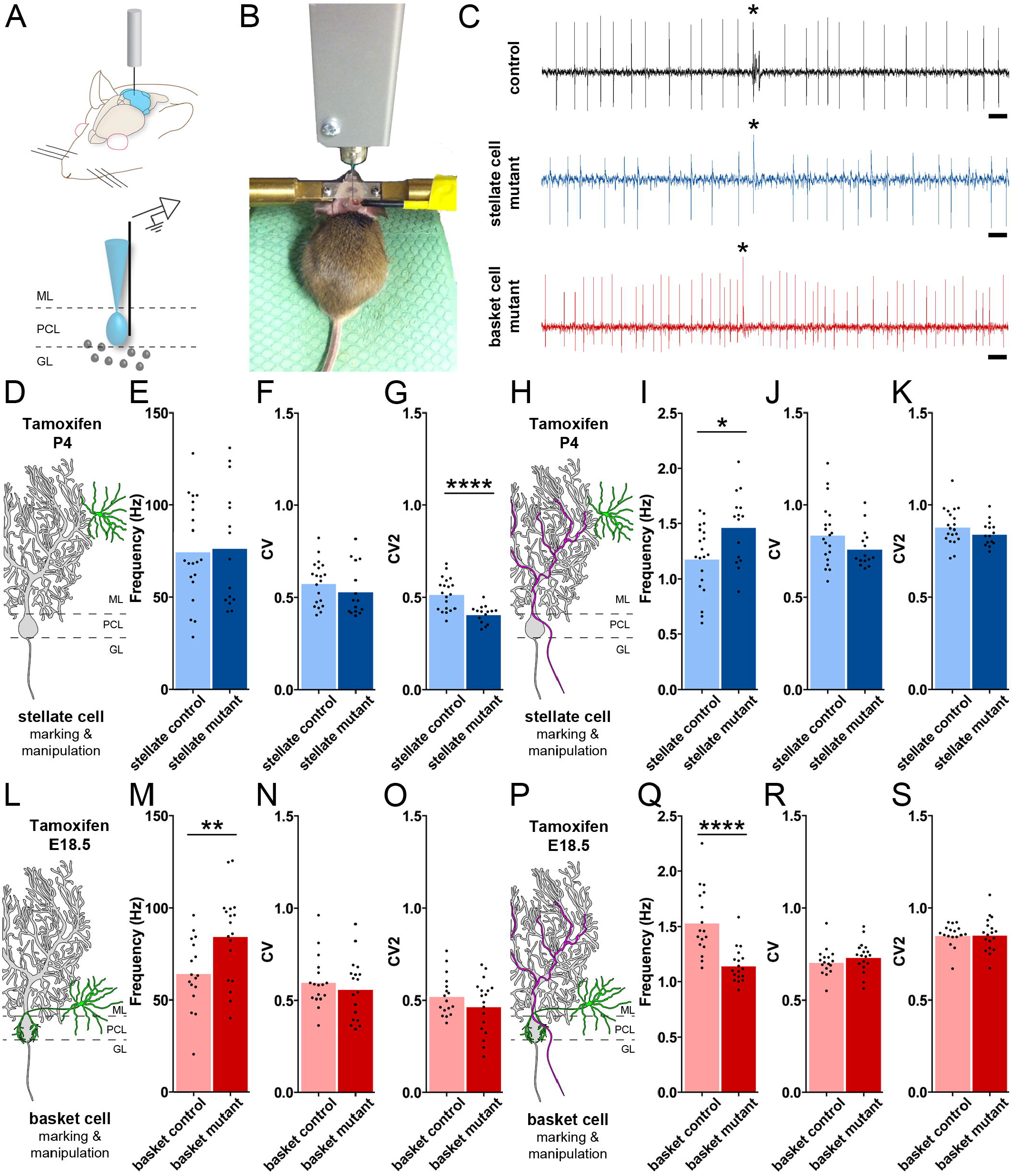
Genetic depletion of GABAergic molecular layer interneuron neurotransmission alters Purkinje cell firing *in vivo*. (**A**) Schematic of electrophysiology setup for *in vivo* extracellular recordings. A sharp metal electrode is lowered into the cerebellum of awake mice (above) to target Purkinje cells for single-unit recordings (below). (**B**) Picture of a mouse in the electrophysiology setup. The mouse is headfixed and able to walk on a cylindrical foam wheel. (**C**) Example extracellular recordings of Purkinje cells in a control (top), stellate cell mutant (middle), and basket cell mutant (bottom) mouse. Complex spikes are indicated with asterisks. Scale = 20ms. (**D**) Schematic of a stellate cell (green) in relation to a Purkinje cell (grey). (**E-G**) Quantification of Purkinje simple spike electrophysiology in awake stellate cell control (N = 7, n = 20) and stellate cell mutant (N = 3, n = 15) mice. (E) Quantification of Purkinje cell simple spike firing frequency in stellate cell control and mutant conditions. Firing frequency was unchanged in the stellate cell silencing condition (stellate cell control = 74.99Hz ± 6.031Hz; VGAT KO = 76.85Hz ± 8.298Hz; *P* = 0.8577). (**F**) Quantification of Purkinje cell simple spike coefficient of variance (CV) in stellate cell control and mutant conditions. CV was not significantly changed from control in the stellate cell mutant animals (stellate cell control mean = 0.5787 ± 0.0264; stellate cell mutant mean = 0.5333 ± 0.03603; *P* = 0.3181.). (**G**) Quantification of Purkinje cell simple spike CV2 in stellate cell control and mutant conditions. CV2 was significantly decreased from control in the stellate cell mutant condition (stellate cell control mean = 0.5186 ± 0.01968; stellate cell mutant mean = 0.4091 ± 0.01220; *P* < 0.0001). (**H**) Schematic of a climbing fiber (magenta) to a stellate cell (green) and a Purkinje cell (grey). (**I-K**) Quantification of Purkinje complex spike electrophysiology in awake stellate cell control (N = 7, n = 20) and stellate cell mutant (N = 3, n = 15) mice. (**I**) Quantification of Purkinje cell complex spike firing frequency in stellate cell control and mutant conditions. Firing frequency was significantly increased over control in the stellate cell silencing condition (stellate cell control mean = 1.186Hz ± 0.06845Hz; stellate cell mutant mean = 1.469Hz ± 0.08279Hz; *P* = 0.0131). (**N**) Quantification of Purkinje cell complex spike coefficient of variance (CV) in stellate cell control and mutant conditions. CV was not significantly changed from control in stellate cell mutant animals (stellate cell control mean = 0.8412 ± 0.03723; stellate cell mutant mean = 0.7650 ± 0.02704; *P* = 0.1076). (**O**) Quantification of Purkinje cell complex spike CV2 in stellate cell control and mutant conditions. CV2 was not significantly changed from control in the stellate cell VGAT deletion condition (stellate cell control = 0.8819 ± 0.02203; stellate VGAT KO = 0.8456 ± 0.01697; *P* = 0. 2006). (**L**) Schematic of a basket cell (green) in relation to a Purkinje cell (grey). (**M-O**) Quantification of Purkinje simple spike electrophysiology in awake basket cell control (N = 5, n =17) and basket cell mutant (N = 3, n = 18) mice. (**M**) Quantification of Purkinje cell simple spike firing frequency in basket cell control and mutant conditions. Firing frequency was significantly increased over control in the basket cell silencing condition (basket cell control mean = 64.56Hz ± 4.615Hz; basket cell mutant mean = 84.76 ± 5.670Hz; *P* = 0.0094). (N) Quantification of Purkinje cell simple spike coefficient of variance (CV) in basket cell control and mutant conditions. CV was not significantly changed from control in basket cell mutant animals (basket cell control mean = 0.5996 ± 0.03410; basket cell mutant mean = 0.5611 ± 0.03852; *P* = 0.4599). (**O**) Quantification of Purkinje cell simple spike CV2 in basket cell control and mutant conditions. CV2 was not significantly changed from control in the basket cell VGAT deletion condition (basket cell control = 0.5223 ± 0.02673; basket cell VGAT KO = 0.4688 ± 0.03437; *P* = 0.2282). (**P**) Schematic of a climbing fiber (magenta) to a basket cell (green) and a Purkinje cell (grey). (Q-S) Quantification of Purkinje complex spike electrophysiology in awake basket cell control (N = 5, n = 17) and basket cell mutant (N = 3, n = 18) mice. (**Q**) Quantification of Purkinje cell complex spike firing frequency in basket cell control and mutant conditions. Firing frequency was significantly decreased from control in the basket cell silencing condition (basket cell control mean = 1.538Hz ± 0.07298Hz; basket cell mutant mean = 1.148Hz ± 0.03695Hz; *P* < 0.0001). (**R**) Quantification of Purkinje cell complex spike coefficient of variance (CV) in basket cell control and mutant conditions. CV was not significantly changed from control in basket cell mutant animals (basket cell control mean = 0.7074 ± 0.01833; basket cell mutant mean = 0.7342 ± 0.01970; *P* = 0.3252). (**S**) Quantification of Purkinje cell complex spike CV2 in basket cell control and mutant conditions. CV2 was not significantly changed from control in the basket cell VGAT deletion condition (basket cell control = 0.8533 ± 0.01476; basket VGAT KO = 0.8552 ± 0.02171; *P* = 0.9414).

### Loss of molecular layer interneuron inhibition does not cause neurodegeneration

We wondered whether removing GABAergic neurotransmission from molecular layer interneurons altered Purkinje cell function because of neurodegeneration in the cerebellum. This was important to test because dysgenesis during development and neurodegeneration in the adult are known to drive a number of electrophysiological abnormalities (Reeber et al., 2013). Specifically, alterations of the Purkinje cell dendrites after loss of interneuron connectivity would be a primary concern in our paradigm. We therefore measured molecular layer thickness as a proxy for dendrite span. Molecular layer thickness is a sensitive and straightforward measure for developmental and adult-associated defects that disrupt Purkinje cell dendrite size (Hansen et al., 2013; White and Sillitoe, 2017; White et al., 2014, 2016b). We stained sagittal cut tissue sections of the cerebellum with either anti-calbindin or anti-CAR8 antibody, which mark Purkinje cells, and a fluorescent Nissl stain or DAPI, which outline all layers but heavily mark the granular layer because of the high cell density (Figure 4A-B). Molecular layer thickness was assessed for lobule III/IV by measuring the perpendicular distance from the molecular layer-facing edge of a Purkinje cell soma to the outer edge of the molecular layer. We found that molecular layer thickness was not altered in either of the mutant mice compared to controls (Figure 4C; stellate cells – control = 159.7μm ± 9.201; mutant = 157.2μm ± 5.493; *P* = 0.8288; basket cells – control = 179.9μm ± 3.833; mutant = 181.1μm ± 3.164; *P* = 0.8200). These data indicate that the outgrowth of the Purkinje cell dendritic tree during postnatal development, and its maintenance thereafter, were not adversely affected after we genetically silenced stellate cell and basket cell GABAergic output activity in the developing cerebellar cortex.

**Figure 4.**
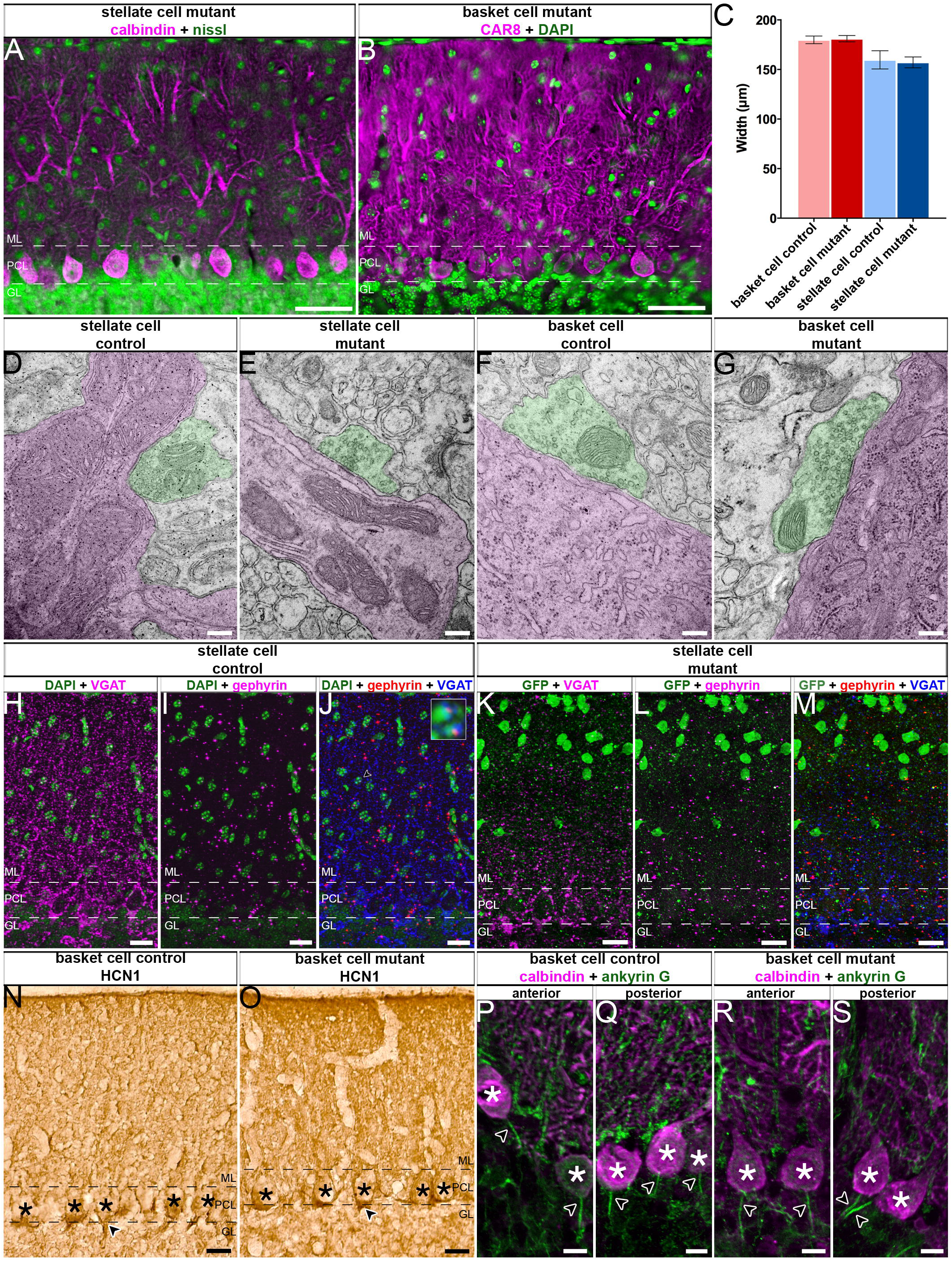
Deleting *Vgat* in molecular layer interneurons does not rearrange cerebellar circuitry or induce neurodegeneration. (**A-B**) Example images of sagittal cerebellar sections used for quantification of ML thickness wherein Purkinje cells were stained with either calbindin or CAR8 and all neurons were stained with either nissl or DAPI to facilitate visibility of the Purkinje cell layer (PCL) compared to the molecular layer (ML) and granule layer (GL). Scale = 50μm. ML thickness was unchanged in the stellate silencing condition (**A**) or the basket cell silencing condition (**B**). (**C**) Quantification of ML thickness in all conditions. ML thickness is not significantly changed from control in either basket cell or stellate cell mutant animals (basket cell control mean = 179.9μm ± 3.833, basket cell mutant mean = 181.1μm ± 3.164, *P* = 0.8200; stellate cell control mean = 159.7μm ± 9.201, stellate cell mutant mean = 157.2μm ± 5.493, *P* = 0.8288). (**D-G**) TEM images revealed normal synapses in all conditions. Purkinje cells and processes are colorized in magenta and identified basket and stellate synaptic terminals are colorized in green. Scale = 200nm. Inhibitory synapses onto Purkinje cell dendrites in the ML were unchanged from the control (**D**) in stellate cell mutant mice (**E**). Similarly, inhibitory synapses onto Purkinje cell somas were unchanged from the control (**F**) in basket cell mutant mice (**G**). (**H-M**) Gephyrin expression was unchanged in stellate cell mutant mice compared to control. Scale = 20μm. Control mice (**H-J**) have uniform expression of VGAT in the ML (**H**) and similarly uniform expression of gephyrin at inhibitory synapses in the ML (**I**). Triple staining reveals gephyrin is present at inhibitory synapses in the ML. Example triple labeled synapses (arrowhead) are shown in the blowup (**J**). Stellate cell mutant mice do not have uniform expression of VGAT in the ML as a result of the targeted deletion of VGAT (**K**). However, gephyrin appears uniformly expressed (L) suggesting it is present at synapses as normal, despite the depletion of VGAT (**M**). (**N-S**) Postsynaptic structures are also unchanged in basket cell mutant mice. HCN1 staining suggests the region of the basket cell pinceau is unchanged from control (**N**) in basket cell mutant mice (**O**). Scale = 20μm. (**P-S**) The Purkinje cell axon initial segment (stained with ankyrin G and indicated by arrowheads) is obvious in control (**P-Q**) and basket cell mutant mice (**R-S**) throughout the cerebellum with example images show from both anterior and posterior lobules. Purkinje cells are stained with calbindin with their somas indicated by asterisks. Scale = 10μm. (**A-B, H-S**) Dotted lines indicate the borders of the Purkinje cell layer (PCL) with the molecular layer (ML) above and the granule layer (GL) below.

### Deleting *Vgat* in interneurons does not alter their targeting onto Purkinje cells

We next wanted to determine whether interneurons that lack *Vgat* are targeted to the correct regions of the Purkinje cell. We therefore examined whether the ultrastructure of synapses in the molecular layer was intact. To do so, we performed electron microscopy on sagittal sections cut through the adult cerebellum. Using the distinctively large soma of Purkinje cells as a reference point for where the molecular layer starts, we assessed the integrity of inhibitory synapses in the Purkinje cell layer and molecular layer. Stellate cells terminate on the shaft of the Purkinje cell dendritic tree (Palay and Chan-Palay, 1974). Excitatory synapses are distinguished from inhibitory synapses by the presence or absence, respectively, of a postsynaptic density that gives excitatory synapses an asymmetric appearance (Palay and Chan-Palay, 1974). We observed synapses with symmetric morphologies that form postsynaptic terminals on the Purkinje cell dendrites in the molecular layer (Figure 4D-E). These findings indicate that inhibitory synapses are retained in their correct positions within the cerebellar cortex despite the conditional silencing of stellate cell interneuron synapses. We performed a similar analysis in mice with silenced basket cell output. The axons of several basket cells converge on single Purkinje cell somata to form the basket (Palay and Chan-Palay, 1974). Basket cell axons extend further to form specialized pinceaux synapses around the axon initial segments of Purkinje cells (Ango et al., 2004; Palay and Chan-Palay, 1974; Sotelo, 2008). We found inhibitory synapses on the Purkinje cell somata (Figure 4F-G). Importantly, the interneuron synapses in the stellate cell and basket cell mutants contain distinct vesicles. This result indicates that despite the deletion of *Vgat* and the loss of GABAergic neurotransmission, the synaptic structural machinery that is required for housing neurotransmitters before release, remains intact (Figure 4D-G).

To complement the electron microscopy studies in which we assessed presynaptic components, we also tested the correct distribution of the postsynaptic structures belonging to the inhibitory synapses by immunohistochemical staining and light microscopy. Gephyrin is expressed in the postsynaptic compartment of inhibitory synapses (Sassoè-Pognetto et al., 1999). In *Ascl1^CreER/+^; R26^fx-stop-EYFP^;Vgat^fx/fx^* mutant mice treated with tamoxifen at P4, triple staining with gephyrin, VGAT, and GFP revealed a normal distribution of gephyrin in GFP-rich molecular layer regions that were devoid of VGAT expression (Figure 4H-M). After silencing basket cells by giving tamoxifen at E18.5 to *Ascl1^CreER/+^; Vgat^fx/fx^* mutants, we found that HCN1 (hyperpolarization-activated cyclic nucleotide-gated channel), which is expressed at both the pre-and post-synaptic sites at basket cell to Purkinje cell connections (Luján et al., 2005), had a normal expression profile around the Purkinje cell layer (Figure 4N-O). Moreover, we used AnkG (ankyrin-G) expression to show the presence of Purkinje cell axon initial segments after the loss of basket cell inhibitory neurotransmission in *Ascl1^CreER/+^; Vgat^fx/fx^* mutant mice (Figure 4P-S; Buttermore et al., 2012). We also sought to determine whether other major cell types of the cerebellar cortex were present as normal, since abnormal Purkinje cell activity may affect the gross organization of cerebellar circuitry. Purkinje cells, granule cells, Golgi cells, parallel fibers, mossy fibers, climbing fibers, and unipolar brush cells were all present with similar location and morphology in both the basket and stellate cell *Vgat* mutants as compared to control cerebella (Figure 5). Finally, we sought to determine which *Ascl1* lineage cells had been manipulated outside of the cerebellum at both time points to determine whether it was likely the deletion of *Vgat* from these cells could result in the alterations in Purkinje cell firing that were found. Similar to previous work (Kim et al., 2008), we found the majority of *Ascl1* lineage extracerebellar cells differentiating at both our basket and stellate time points are oligodendrocytes and olfactory bulb neurons (Figure 6). We therefore predicted that in our *Vgat* deletion paradigms, relatively few cells would have been manipulated outside of the cerebellum (Figure 6A & H). Specifically, based on reporter expression we found that the majority of extracerebellar cells outside the olfactory bulb had a glial-like morphology (Figure 6B-G, I-N). These putative glial cells were co-labeled with carbonic anhydrase II (CAII), suggesting their identity as oligodendrocytes (Figure 6O). The extracerebellar cells with a more neuron-like morphology included very sparse putative granule cells in the hippocampus that were detected only in the stellate cell marking scheme (Figure 6K) and olfactory bulb neurons that were detected in both the stellate and basket cell marking schemes (Figure 6G). The identity of these cells as neurons was confirmed by the co-labeling of GFP reporter and NeuN (Figure 6P-Q). The vast majority of the non-glial extracerebellar cells were found in the olfactory bulb. These results indicate that the extracerebellar deletion of *Vgat* occurred in a population consisting largely of oligodendrocytes and olfactory bulb neurons, a population of neurons from which the deletion of *Vgat* gene function would be highly unlikely to have significant effects on cerebellar Purkinje cell activity.

**Figure 5.**
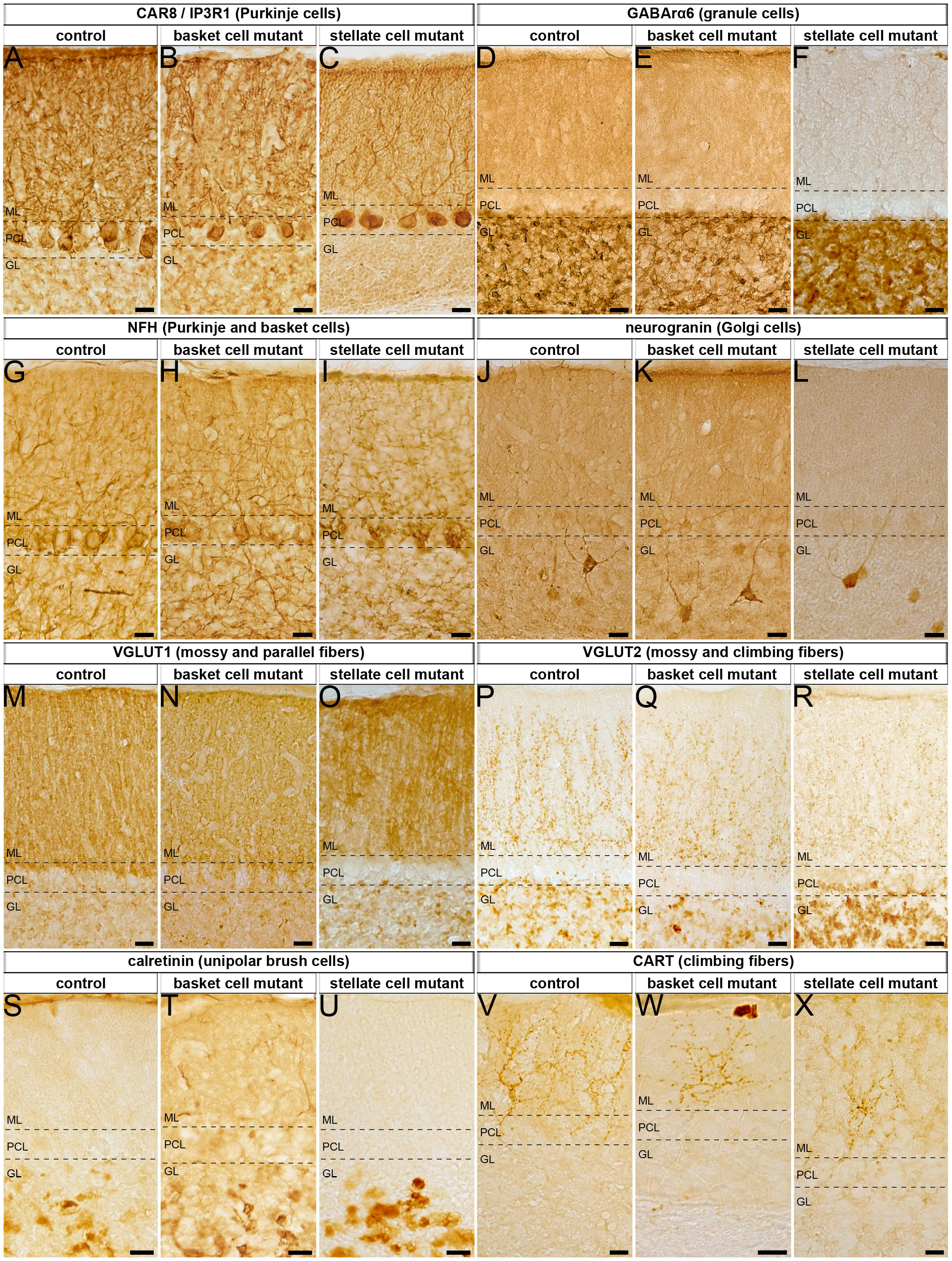
Conditional deletion of *Vgat* in molecular layer interneurons does not lead to gross cerebellar changes in cellular composition, cellular distribution, or layer patterning. (**A-X**) Cerebellar cell types were present and appeared unchanged in location and morphology despite the lack of VGAT in basket cells and stellate cells. Dotted lines indicate the borders of the Purkinje cell layer (PCL) with the molecular layer (ML) and the granule layer (GL). Scale = 20μm. CAR8 and IP3R1 staining revealed normal Purkinje cell location and morphology (**A-C**). GABAαR6 showed normal expression in granule cells (**D-F**). NFH expression was unchanged in Purkinje and basket cells (**G-I**). Neurogranin expression in Golgi cells was unchanged in all conditions (**J-L**). In the mutants, VGLUT1 staining in mossy and parallel fibers was unchanged (**M-O**) compared to control conditions and similarly VGLUT2 was present in mossy fiber terminals in the granule cell layer and climbing fibers in the molecular layer (**P-R**). Staining of unipolar brush cells with calretinin was similar between controls and mutants (**S-U**). CART staining of climbing fibers in the mutants was also consistent with controls (**V-X**).

**Figure 6.**
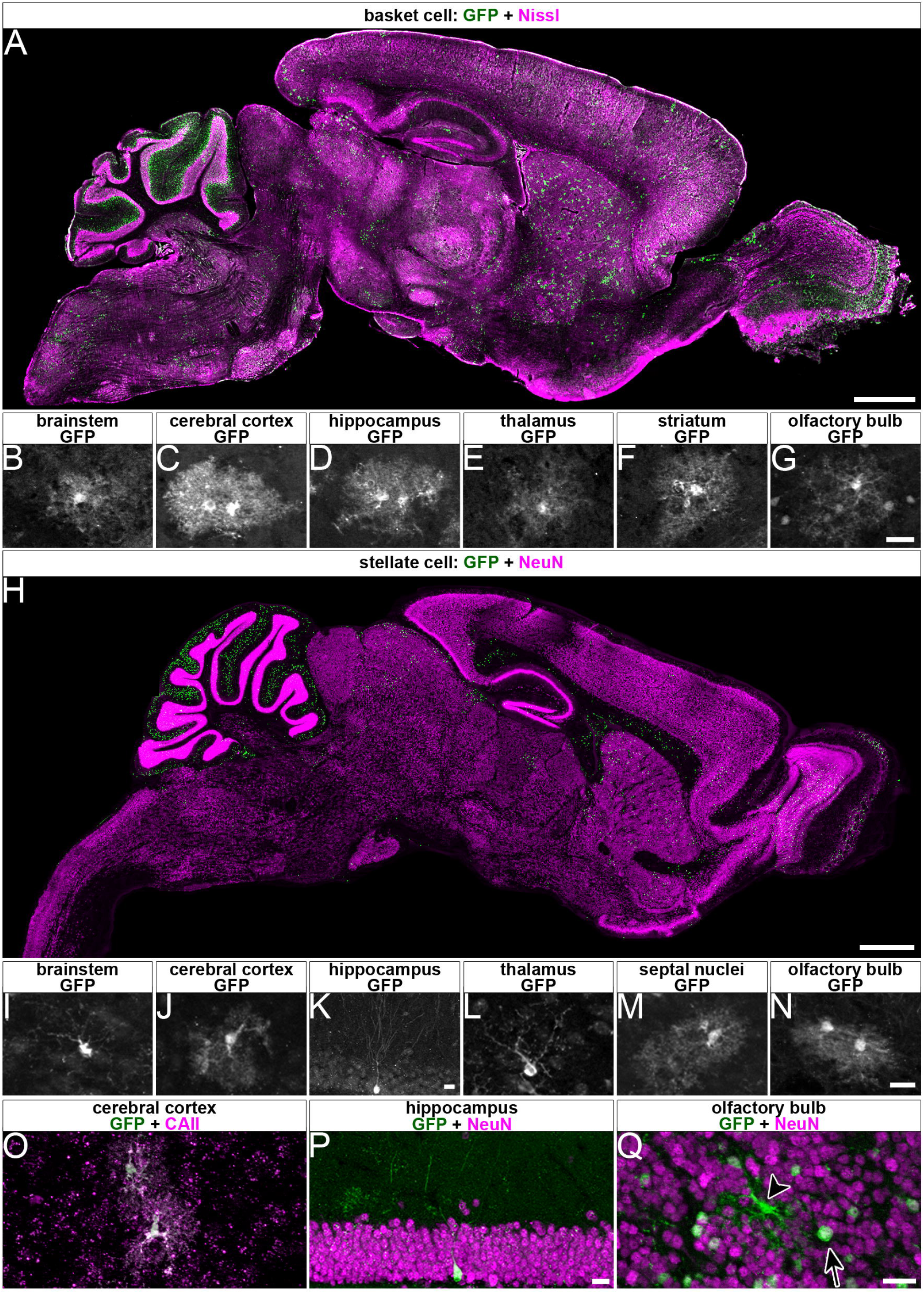
Conditional deletion of *Vgat* with *Ascl1^CreER^* occurs in extracerebellar cell types that are unlikely to affect Purkinje cell activity in this manipulation. (**A**) Sparse labeling of cells occurs outside the cerebellum at the basket cell time point. Scale = 1mm. (**B-G**) Many of the cells outside the cerebellum had morphologies that resembled glia, with the notable exception of cells in the olfactory bulb (**G**) where the majority of cells had the morphology of neurons, though cells with glial-like morphology were also present. Scale = 20μm. (**H**) Sparse labeling outside of the cerebellum also occurred at the stellate time point. Scale = 1mm. (**I-N**) While again many of the cells had morphologies that resembled glia, some cells with neuron-like morphologies were also detected. (**K**) Very sparse labeling of putative granule cells in the dentate gyrus occurred at the stellate time point, unlike at the basket cell time point at which no neurons were detected in the hippocampus. (**N**) Similar to the basket cell time point, many neurons in the olfactory bulb were labeled in addition to some glial-like cells. Scale = 20μm. (**O**) Recombined cells with glial-like morphologies co-labeled with GFP and CAII, a maker of oligodendrocytes. Scale = 20μm. (**P**) Cells with neuron-like morphologies in the dentate gyrus of the hippocampus colabeled with GFP and NeuN. Scale = 20μm. (**Q**) Both neurons (arrow, co-labeled with GFP and NeuN) and glia (arrowhead, only labeled with GFP and not by NeuN) were labeled in the olfactory bulb. Scale = 20μm.

## Discussion

The cerebellum has served as the structure of choice in thousands of developmental, anatomical, functional, and behavioral studies. Among the reasons for its popularity are that its main cell types were identified more than a century ago (Ramón y Cajal, 1909), and electrophysiological methods have allowed a detailed understanding of its connections (Eccles et al., 1976). However, it is still unclear how connectivity within the different classes of interneurons influences cerebellar cortical function. This study is focused on understanding whether the molecular layer interneurons have distinct inhibitory impacts on their target Purkinje cells. We tested how stellate cell and basket cell GABAergic neurotransmission influences Purkinje cell activity. To address this problem, we devised a genetic approach in which we used an *Ascl1^CreER^* mouse line to delete the *Vgat* gene in the developing cerebellum. The *Ascl1^CreER^* allele provided an opportunity for spatial and temporal manipulation of stellate cells independently from basket cells (Sudarov et al., 2011). We found that loss of *Vgat* in stellate cells altered the pattern of Purkinje cell simple spike firing and the rate of complex spike firing in alert mice, whereas deleting *Vgat* in basket cells changed the rate of both Purkinje cell simple spike and complex spike firing. The data suggest that molecular layer interneurons cooperate to establish Purkinje cell function *in vivo.*

### Are cerebellar stellate cells and basket cells distinct cell types?

Traditional high-resolution anatomy distinguishes molecular layer inhibitory interneurons based on multiple cellular, sub-cellular, and connectivity features (Palay and Chan-Palay, 1974). Still, even using these various features it can be difficult to unambiguously assign neurons to a specific stellate or basket cell identity. Golgi staining analysis later suggested that classification based on distinct groups is challenging at best, since a more gradual and continuous identity could better reflect the molecular layer composition (Sultan and Bower, 1998). Analysis of gene expression yet again challenged the view, as the differential expression of multiple genes indicates at least some level of specificity and potentially unique identities within the interneurons (Schilling and Oberdick, 2009). Despite the differential expression, the authors also argue for a common origin. Indeed, stellate cells and baskets arise from a common precursor pool in the ventricular zone (Hoshino et al., 2005), and they are generated in waves during embryonic through postnatal development (Leto et al., 2009; Sudarov et al., 2011). These different perspectives are further complicated by the observation that even though the somata are located in distinct positions within the dorsal-ventral axis of the molecular layer, there is some spread of both cell types’ somata into the middle molecular layer and a fuzzy separation of synaptic location (Figure 2B and 2G). Regardless of anatomical or molecular distinctness, we asked whether any of these properties impact their contribution to cerebellar function. There is consensus that stellate cells and basket cells both synapse directly onto Purkinje cells (Palay and Chan-Palay, 1974). But, do they influence Purkinje cells in a similar or different manner? To tackle this question, we used an *in vivo* genetic model in which fast GABAergic neurotransmission is blocked without causing neurodegeneration or overt circuit rearrangements that would, if present, alter Purkinje cell function. Genetic deletion of VGAT, in general, does not impair the development of inhibitory synapses (Wojcik et al., 2006). Nor does it alter the gross morphology or the basic structure of cerebellar circuits (White et al., 2014). Our results uncover that stellate cells and basket cells do have distinct functional interactions with their Purkinje cell targets, with stellate silencing influencing Purkinje cell simple spike pattern and complex spike rate (Figure 3G and 3I) and basket cell silencing altering the rate of both simple and complex spikes (Figure 3N and 3Q). However, our data cannot exclude the possibility that both cell types modulate multiple aspects of Purkinje cell function, even though each one might have a preferred interaction for modulating rate compared to pattern. In other words, there is likely no one molecular layer inhibitory cell type dedicated exclusively for control of rate and pattern. In slice, inhibitory activity was shown to control the regularity of interneuron firing (Häusser and Clark, 1997), and in a specific form of inhibitory rebound plasticity, basket cells were shown to control the pattern and rate of Purkinje cell output (He et al., 2015). It would be interesting if basket cells and stellate cells are co-opted for rate versus pattern modulation depending on the specific behavioral task or the specific changes in plasticity that arise. Indeed, based on our current data recorded *in vivo*, we can speculate that the predominant roles of the two classes of interneuron might be strengthened by network activity at the population level. Given the developmental nature of our manipulation, it is also possible the consequences we observed on Purkinje cell firing are due, at least in part, to compensatory or plasticity mechanisms after *Vgat* deletion. Even if this were the case, it is still intriguing that Purkinje cell rate is refractory to loss of stellate cell input whereas pattern is refractory to basket cell input.

Connectivity within molecular layer interneurons might be organized in a manner that harnesses their unique developmental properties, wiring diagrams, and functional roles Stellate cells and basket cells do not function in isolation, and interactions within each cell type are not entirely random. Rather, the electrical and chemical connectivity in molecular layer interneuron populations are both highly structured, with connectivity clustering coefficients that reflect a spatial arrangement in the sagittal plane (Rieubland et al., 2014). This architecture is intriguing because the entire cerebellum is organized around a map of sagittal compartments (Apps and Hawkes, 2009; Cerminara et al., 2015). With specific importance to molecular layer interneuron circuitry, it is the Purkinje cells that determine all aspects of cerebellar sagittal organization. Purkinje cells are organized into a complex but precisely pleated array of sagittal compartments that are defined by cellular birth dates, lineage, gene expression, afferent connectivity, and neuronal firing properties (Cerminara et al., 2015; White and Sillitoe, 2013b). Purkinje cells cues during development establish the fundamental map (Croci et al., 2006; Sillitoe et al., 2008a) whereas Purkinje cells activity fine-tunes the topography into functional modules (White et al., 2014). Molecular markers link subsets of interneurons to specific Purkinje cells forming zones defined by common expression (Chan-Palay et al., 1982). There is also some evidence that the inhibitory neurons follow the expression of zebrinII (Sillitoe et al., 2008b), the most extensively studied molecular marker of Purkinje cell zones (Brochu et al., 1990; Sillitoe and Hawkes, 2002). Based on the *Ascl1^CreER^* marking schemes for stellate cells and basket cells, there is no reason to believe that either paradigm marked cells that were restricted to particular zonal compartments (Figure 2F-G), although it is possible that an interneuron’s birth date determines the particular zonal circuit that it will eventually wire into.

Deletion of *Vgat* using *Ascl1^CreER^* was predicted to leave signaling intact in a substantial number of cells. By design, only subpopulations of molecular layer interneurons were targeted, resulting in total molecular layer recombination of ~35% for each scheme (Figure 2E) with the majority of labeled cells found in their canonical regions of the molecular layer (Figure 1F-G). Still, VGAT was not entirely eliminated, but was instead reduced by ~57% in the basal molecular layer and ~64% in the apical molecular in the basket and stellate schemes, respectively (Figure 2I-J). This efficiency is impressive for only a single dose of tamoxifen given that the molecular layer interneurons are born progressively over several embryonic and postnatal days. Still, even by creating a mosaic population of silenced interneurons we detected significant deficits in the overall function of Purkinje cells regardless of which particular cerebellar zone the recorded cell resided within (Xiao et al., 2014; Zhou et al., 2014). Part of the reliability in producing Purkinje cells firing defects could be due to the connectivity of each manipulated interneuron, given that each one has the potential to make synaptic contacts with multiple Purkinje cells (Palay and Chan-Palay, 1974). While stellate cells make mainly local synaptic connections with potentially fewer long-distance contacts, the basket cells could contact upwards of 9 Purkinje cells each (Palkovits et al., 1971). The establishment of these distributions could also be altered in our genetic deletion paradigms. During development, synaptic activity controls the speed and direction of migration (Wefers et al., 2017). Because stellate cells and basket cells have intra-and inter-cellular connections with one another (Palay and Chan-Palay, 1974), loss of GABAergic neurotransmission could impede neuronal migration. We suspect that if there were such deficits, they would likely be subtle or highly localized and specific since we did not detect obvious changes in cerebellar cell distribution by immunohistochemistry (Figure 2C, 4A-B, 5A-X) or afferent targeting as determined by electron microscopy (Figure 4D-G).

### Implications of interneuron connectivity on cerebellar circuit function

Despite a long and rich history of understanding cerebellar cellular composition, circuitry, and function (Eccles, 1967; Ramón y Cajal, 1909; Voogd, 2014; Voogd and Glickstein, 1998), the last decade of cerebellar research has uncovered a number of additional cerebellar cortical afferent and efferent connections that could influence molecular layer interneuron processing. Purkinje cells not only contact the cerebellar nuclei, but through collaterals they also contact each other (Díaz-Rojas et al., 2015; Orduz and Llano, 2007; Orduz et al., 2014; Watt et al., 2009; Witter et al., 2016), interneurons (Witter et al., 2016), and granule cells (Guo et al., 2016). The cerebellar nuclei indeed project out of the cerebellum, but they too also project back to the cerebellar cortex by way of inhibitory processes to Golgi cells and excitatory processes to Golgi cells (Ankri et al., 2015) and granule cells (Gao et al., 2016; Houck and Person, 2015). In this context, we should consider the various possible inputs to the molecular layer interneurons: climbing fibers to stellate cells and basket cells, Purkinje cells to stellate and basket cells, granule cells to stellate and basket cells, stellate cells to basket cells, basket cells to stellate cells, basket cells to basket cells, and stellate cells to stellate cells (Palay and Chan-Palay, 1974; Witter et al., 2016). Before each interneuron communicates its output to its respective Purkinje cells, we also take into account that electrical connections tether rodent basket cells into groups of 5 and stellate cells in pairs (Alcami and Marty, 2013). Moreover, the interaction between small patches of granule cells and Purkinje cells is shaped by molecular layer interneurons, and the strength of this inter-layer communication is dependent on relative position to the Purkinje cells in the sagittal and mediolateral axis (Dizon and Khodakhah, 2011). It should also be considered that although the molecular layer interneurons are defined as GABAergic, they exhibit the expected inhibitory drive as well as a less appreciated excitatory influence (Chavas and Marty, 2003). Specifically, for our stellate cell silencing paradigm, it could be that the lack of a change in simple spike rate indicates an equilibrium rather than the absence of an effect. The predicted increase in Purkinje cell firing rate after loss of inhibitory GABA function would be countered by decrease in Purkinje cell spikes after removing excitatory GABA function (Figure 3). Under normal physiological conditions, such an effect could have a modulatory role in finely controlling Purkinje cell spike output, especially when dynamic changes are required during unrestricted behavior (Sauerbrei et al., 2015). The impact of interneuron communication perhaps could also be appreciated at the population level. It could be that the local electrical networking together with their arrangement into rows facilitates a topographic interaction with zonally projecting climbing fibers from the inferior olive (Lang et al., 2017; Sugihara et al., 2009). At the level of Purkinje cells, this ordered cellular and circuit architecture could manifest as synchronous activity (Lang et al., 2014). Synchrony between chemically linked molecular layer interneurons has been reported (Rieubland et al., 2014) and their impact is likely restricted to sagittal bands (Mann-Metzer and Yarom, 1999). This is consistent with the long-standing hypothesis that synchronous neural activity promotes a level of neuronal ensemble dynamics that allow for muscles synergies to accommodate complex motor behaviors (Welsh et al., 1995).

## Conclusions

Cerebellar stellate cells and basket cells are the predominant cell type of the molecular layer. They arise from a common progenitor pool in the ventricular zone of the cerebellum and continue to divide and differentiate through postnatal development. We used an *Ascl1^CreER^* genetic inducible allele to leverage this spatial and temporal pattern of development in order to manipulate the synaptic output of inhibitory interneurons. By blocking *Vgat* expression and then recording Purkinje cell activity in alert adult mice we uncovered that stellate cells establish the Purkinje cell simple spike firing pattern whereas basket cells determine their rate. Additionally, we found that Purkinje cell complex spike firing rate increases with a lack of stellate cell inhibition but in contrast decreases with a lack of basket cell inhibition. This study establishes complementary roles for the GABAergic function of cerebellar molecular layer interneurons.

## Materials and Methods

### Mouse Lines

All experiments were performed according to a protocol approved by the Institutional Animal Care and Use Committee (IACUC) of Baylor College of Medicine. Three mouse lines were intercrossed to generate the various alleles. The first line expresses a knock-in construct of the *CreER^T2^* allele under the control of the *Ascl1* promoter *(Ascl1^CreER^)* (Sudarov et al., 2011). The second line carries a knock-in floxed *Vgat* allele *(Vgat^fx^*) (Tong et al., 2008). The third line expresses an enhanced yellow fluorescent protein (EYFP) knock-in construct with an upstream floxed transcriptional stop sequence, under the control of the *ROSA26* locus *(R26^fx-stop-EYFP^*) (Srinivas et al., 2001). Our genotyping procedures for all of these alleles have been described before (Sillitoe et al., 2009; White and Sillitoe, 2017; White et al., 2014). We bred the mice using standard timed pregnancies, and we designated noon on the day a vaginal plug was detected as embryonic day (E) 0.5 and the day of birth as P0. Mice of both sexes were studied. The mice were housed on a 14h/10h light/dark cycle.

### Cre induction

Tamoxifen (Sigma) was dissolved at 37°C overnight in corn oil at a concentration of 20 mg/ml (Sillitoe et al., 2009; Zervas et al., 2004). An 18-gauge syringe was used to pipette the solution up and down and dissolve any remaining tamoxifen particles. For targeting the stellate cells, tamoxifen was delivered at a dosage of 200ug/g into P4 postnatal pups by subcutaneous injection into the skinfold at the back of the neck. The pups were allowed to rest in a separate cage to prevent the mother from licking out the tamoxifen. After ~15 minutes, or once the subcutaneous bolus of tamoxifen solution had completely dispersed, each pup was returned to its home cage. For targeting the basket cells, 200ug/g tamoxifen was add-mixed with 50ug/g progesterone and administered to pregnant dams by oral gavage (Bowers et al., 2012).

### Immunohistochemistry

Perfusion and tissue fixation were performed as previously described (Sillitoe et al., 2008a). Briefly, mice were anesthetized by intraperitoneal injection with Avertin (2, 2, 2-Tribromoethanol, Sigma-Aldrich Cat # T4). Cardiac perfusion was performed with 0.1 M phosphate-buffered saline (PBS; pH 7.4), then by 4% paraformaldehyde (4% PFA) diluted in PBS. For cryoembedding, brains were post-fixed in 4°C for 24 to 48 hours in 4% PFA and then cryoprotected stepwise in sucrose solutions (15% and 30% diluted in PBS) and embedded in Tissue-Tek^®^ O.C.T. Compound (Sakura, Torrance, CA, USA). Samples were cut on a cryostat with a thickness of 40 μm and sections were collected as free-floating sections and stored in PBS. Immunohistochemistry procedures on free-floating frozen tissue sections were described previously (Sillitoe et al., 2003, 2010; White and Sillitoe, 2013a; White et al., 2014). After staining, the tissue sections were placed on electrostatically coated slides and allowed to dry.

### Cerebellar circuit markers

The integrity of the cerebellar circuitry was checked by determining the expression patterns of several synaptic and cell type-specific markers. Excitatory glutamatergic synapses contributed by granule cell parallel fibers were immunolabeled with rabbit anti-vesicular glutamate transporter 1 (anti-VGLUT1; 1:1000; Synaptic Systems, Göttingen, Germany). Excitatory synapses contributed by the mossy fibers in the granular layer (Gebre et al., 2012) and the climbing fibers in the molecular layer (Hisano et al., 2002) were immunolabeled with rabbit anti-VGLUT2 (1:500; Synaptic Systems, Göttingen, Germany; Cat. # 135 403) and rabbit polyclonal anti-cocaine- and amphetamine-related transcript peptide (CART; 1:250; Phoenix Pharmaceuticals, Burlingame, CA, USA; Cat. # H-003-62). The CART signal was amplified using a biotinylated secondary antibody (Vectastain Elite ABC method; Vector Labs; Burlingame, CA, USA) and used to visualize climbing fibers mainly in lobules IX and × (Reeber et al., 2011).

Purkinje cells were marked with anti-calbindin (1:1,000; Cat. # 300; Swant, Marly, Switzerland), rabbit polyclonal anti-carbonic anhydrase or CAR8 (CAVIII, 1:1000l; Cat. # sc-67330, Santa Cruz Biotechnology), goat polyclonal anti-IP3R1 (1:500; Cat. # sc-6093, Santa Cruz Biotechnology, Dallas, TX, USA), goat polyclonal anti-RORα (1:250; Cat. # sc-6062, Santa Cruz Biotechnology, Dallas, TX, USA), and mouse monoclonal anti-ankyrin-G (1:200; Cat. # MABN466, clone N106/36, Millipore Sigma, Burlington, MA, USA). Purkinje cells and molecular layer interneurons were marked with rabbit polyclonal anti-parvalbumin (1:1000; Swant, Marly, Switzerland; Cat. # PV25). Excitatory interneurons were marked by rabbit polyclonal anti-calretinin (1:500; Swant, Marly, Switzerland; Cat. # CR7699/3H). Granule cells were marked with rabbit polyclonal anti-gamma-aminobutyric acid receptor α6 (GABARα6; 1:500; Millipore Sigma, Burlington, MA, USA; Cat. # AB5610). Golgi cell interneurons in the adult cerebellum were marked by rabbit polyclonal anti-neurogranin (1:500; Millipore Sigma, Burlington, MA, USA; Cat. # AB5620) (Singec et al., 2003). NeuN (1:250; Millipore Sigma, Burlington, MA, USA; Cat. #mab377) was used as a general neuronal marker and carbonic anhydrase II (CAII; BioRad, Hercules, CA, USA; Cat. # 00073) was used to label oligodendrocytes. Neuronal processes were also labeled with various markers. Mouse monoclonal anti-neurofilament heavy (NFH; also called anti-SMI-32; 1:1500; Covance, Princeton, NJ) immunolabeled the soma, dendrites, and axons of adult Purkinje cells, and the axons and terminals of basket cells. Mouse monoclonal anti-hyperpolarization-activated cyclic nucleotide-gated channel 1 (HCN1; 1:200; Alomone Labs; Jerusalem, Israel, Cat. # APC-056) was also used to label basket cell axons and pinceaux terminals. Guinea pig anti-gephyrin (1:500; Synaptic Systems, Göttingen, Germany, Cat. #147 004) was processed on paraffin embedded tissue cut at 10 μm. Some tissue sections were double-labeled with the different markers listed above plus chicken anti-GFP (1:1000; Abcam, Cambridge, UK, Cat. # AB13970) in order to visualize the EYFP and mGFP reporter expression.

For fluorescence immunostaining, we used Alexa-488, −555, and −647 secondary goat anti-mouse and anti-rabbit antibodies (1:1500, 1:1500, and 1:1000, respectively; Molecular Probes Inc., Eugene, OR, USA). For chromogenic immunostaining, we used horseradish peroxidase (HRP)-conjugated secondary goat anti-mouse or anti-rabbit antibodies (1:200; DAKO, Carpinteria, CA, USA). Antibody binding was revealed by incubating the tissue in the peroxidase substrate 3,3 ‘ – diaminobenzidine (DAB; Sigma-Aldrich, St Louis, MO, USA), which was made by dissolving a 100 mg DAB tablet in 40 ml PBS and 10 μL 30% H2O2. The DAB reaction was stopped with PBS when the optimal color intensity was reached. To preserve and contrast the fluorescence signal the tissue sections were mounted either with Fluoro-gel (Electron Microscopy Sciences, Hatfield, PA, USA) or a medium containing DAPI (Vectashield Antifade Mounting Medium with DAPI; Cat. # H-1200, Vector Laboratories, Burlingame, CA, USA).

### Imaging of immunostained tissue

Photomicrographs of the tissue sections were captured using Zeiss AxioCam MRm (fluorescence) and AxioCam MRc5 (DAB-reacted tissue sections) cameras mounted on a Zeiss Axio Imager.M2 microscope or on a Zeiss Axio Zoom.V16. Images of tissue sections were acquired and analyzed using either Zeiss AxioVision software (release 4.8) or Zeiss ZEN software (2012 edition). After imaging the tissue, the raw data were imported into Adobe Photoshop CS6 and corrected for brightness and contrast levels. The schematics were drawn in Adobe Illustrator CS6.

### VGAT quantification

We determined whether Cre induction deleted VGAT in interneurons by immunolabeling sagittal tissue sections from 1-month old mice with guinea pig anti-VGAT antibody (1:500; Synaptic Systems, Cat # 131 004; Göttingen, Germany). Images of the molecular layer were acquired with 20x magnification using Zeiss Axioimager microscope, in z-stack and ApoTome mode. Using the Fiji software for analysis, the background was subtracted using the built-in rolling ball method. The same settings were used for control and mutant tissue. The molecular layer was divided dorso-ventrally into three levels, and the levels were saved as regions of interest (ROI). The area and number of puncta in each level was measured using the built-in Analyze Particles function in Fiji and the density of VGAT-positive puncta for each level was calculated. Statistical significance at p < 0.05 was determined using the Student’s *t*-test.

### Molecular layer thickness measurement

Molecular layer thickness was measured from 3 mice per genotype in 3-4 sagittal sections spanning the midline per mouse, with a distance of ~80 μm in between each section. The tissues were immunostained with mouse monoclonal or rabbit polyclonal anti-calbindin (1:1,000; Cat. # 300; Swant, Marly, Switzerland) or anti-carbonic anhydrase to mark the Purkinje cell and molecular layers and NeuroTrace fluorescent Nissl stain (Life Technologies, Grand Island, NY, USA) or DAPI (Vectashield Antifade Mounting Medium with DAPI; Cat. # H-1200, Vector Laboratories, Burlingame, CA, USA) to mark the granular layer. The distance from the edge of the Purkinje cell soma to the apical edge of the molecular layer in the lobule III/IV region was measured using a line measurement tool from Fiji (Schindelin et al., 2012). Measurements for each mouse were averaged and the numbers computed from each genotype were pooled and averaged again to obtain the mean molecular layer thickness. Statistical significance was defined as p < 0.05 using the Student’s t-test.

### Electron Microscopy

Mice were anesthetized with Avertin and perfused with 0.9% room temperature saline, followed by an ice-cold solution of 4% paraformaldehyde and 2% glutaraldehyde in 0.1M sodium cacodylate buffer (pH 7.4; 305-315 mOsm). Brains were harvested and cerebella were sagittally sectioned using a rodent brain matrix while immersed in fixative. The sections were transferred with fixative to a dish and the position of the molecular layer was noted. The region was chopped into pieces measuring less than 1 mm × 1mm. The pieces were aspirated into a sample vial and fixed for 48 hours in 4°C. Samples were treated with 1% Osmium tetroxide in 0.1M cacodylate buffer for secondary fixation, then subsequently dehydrated in ethanol and propylene oxide and embedded in Embed-812 resin (Electron Microscopy Science, Hatfield, PA). Procedures were performed in a Ted Pella Bio Wave microwave oven with vacuum attachment. Tissues were cut with a Leica UC7 microtome into 50nm ultra-thin sections and collected on Formvar-coated copper grids (Electron Microscopy Science, Hatfield, PA). Specimens were then stained with 1% uranyl acetate and 2.5% lead citrate and imaged using a JEOL JEM 1010 transmission electron microscope with an AMT XR-16 mid-mount 16 mega-pixel CCD camera. The images were imported into ImageJ where a smoothing function was applied and then the data were assembled in Adobe Photoshop CS6.

### Surgery

Surgery for awake recordings was performed as detailed in White et al. (White et al., 2016a). Mice were sedated by gas anesthesia using 3% isoflurane, then injected with a ketamine-dexmedetomidine cocktail at a dosage of 80/16 mg/kg, respectively. They were then transferred from the anesthesia chamber to a stereotaxic platform (David Kopf Instruments, Tujunga, CA, USA) and head-fixed with metal ear bars. Sterile surgery techniques were followed. A custom-made headplate was first attached to the Bregma region using Metabond. This headplate was used to affix the mouse’s head to the awake recording apparatus. After the adhesive has dried, a small hole slightly smaller than a 1/16 screw (00-90×1/16 flat point stainless steel machine screws #B002SG89QQ) was drilled to the left of the cerebellar midline. Drilling was stopped before the skull was completely penetrated. An ethanol-sterilized 1/16 screw, which served as an anchor for dental cement, was secured into the drillhole with a screwdriver until it was tightly in place. Another craniotomy was performed on the right side of the midline. A hole approximately ~5 mm in diameter was drilled, taking care not to damage the dura. Once the craniotomy was complete, the hole was covered in triple antibiotic ointment to prepare for the installation of the recording chamber. A piece of straw with a 5-7 mm diameter and a height of 4-5 mm was ethanol-sterilized and air-dried. One end of the straw was dipped in Metabond and carefully placed on top of the craniotomy. Once the adhesive was dry, dental cement (A-M Systems dental cement powder #525000 and solvent #526000) was applied on the outer edge of the straw to fill in holes and to further secure the chamber. After the dental cement had dried, a fresh layer was applied around the straw and the Bregma region where the headplate was attached. After the layer dried, a final layer was applied throughout the site of surgery, including the screw, the top and underside of the headplate, and along the edges of the straw. The skin surrounding the site of the surgery was fixed to the dental cement using 3M Vetbond (#NC0304169) after the cement has completely dried. While the last layer of dental cement was drying, 0.6 mg/kg buprenorphine was injected subcutaneously as an analgesic. After the surgery, the mouse was placed in a warming box (V500, Peco Services Ltd., Cumbria, UK) to prevent hypothermia while the anesthesia wears off. Once the mouse was awake and mobile, it was returned to the home cage. The mouse was allowed to recover for 2-3 days and was given buprenorphine every 6-12 hours. On the third day, training on the running wheel was started. Training sessions were done twice a day for 30 minutes. Before recording, the antibiotic ointment in the chamber was removed using a compressed foam-tipped swab (Cleanfoam^®^ Swab) and replaced with 0.9% w/v NaCl solution. After each recording session, the solution was removed with a cotton tip or by aspiration with a micropipette and fresh antibiotic ointment applied.

### In vivo electrophysiology

Single-unit extracellular recordings were performed as described previously (Arancillo et al., 2015; White and Sillitoe, 2017; White et al., 2016b). During the recordings, the reference electrode tip was immersed in the saline chamber. Tungsten electrodes (Thomas Recording, Giessen, Germany) with an impedance of 5-8 MΩ were controlled from a headstage using a motorized micromanipulator (MP-225; Sutter Instrument Co., Novato, CA, USA). Signals were acquired using an ELC-03XS amplifier (NPI Electronic Instruments, Tamm, Germany) with band-pass filter settings of 0.3-13 kHz. Analog signals were digitized using a CED Power 1401 and stored and analyzed using Spike 2 software (CED, Cambridge, UK).

Purkinje cells were recorded at a depth of approximately 0-2 mm from the tissue surface while the mouse was alert and standing on a wheel. Purkinje cells were identified by the unique presence of two types of action potentials: simple spikes, which are intrinsically generated, and complex spikes, which are generated by climbing fiber input. Neurons from which we obtained clear, continuous recordings lasting 200-300 seconds were included in the analysis. Analysis of firing properties was performed using Spike2, MS Excel, and GraphPad Prism. Firing rate (Hz) was calculated as the number of spikes recorded over a given time period. Coefficient of variance, or CV, was calculated as the ratio of the standard deviation of the interspike intervals (ISI) over the mean ISI. CV2 was calculated with the formula (CV2 = 2|ISIn+1— ISIn|/(ISIn+1+ISIn)), as described previously (Holt et al., 1996). Purkinje cell simple spike and complex spike activity was sorted, analyzed, and data reported as mean ± standard error of the mean (SEM). GraphPad Prism ROUT method of outlier detection was used with Q = 1% to remove outlier cells before further analysis. Statistical analyses were performed with unpaired, two-tailed Student’s t-tests. Statistical significance is indicated in the graphs as *P<0.05, **P<0.01, ***P<0.001, ****P<***0.0001. The number of Purkinje cells that were analyzed for each measurement is indicated with “n”, while the number of mice recorded for each genotype analyzed is indicated with “N”.

## Acknowledgments

This work was supported by funds from Baylor College of Medicine and Texas Children’s Hospital. RVS also received support from the Bachmann-Strauss Dystonia and Parkinson Foundation, Inc., Mrs. Clifford Elder White Graham Endowed Research Fund, the Hamill Foundation, the Caroline Wiess Law Fund for Research in Molecular Medicine, a BCM IDDRC Project Development Award, BCM IDDRC Grant U54HD083092 from the Eunice Kennedy Shriver National Institute of Child Health and Human Development (The IDDRC Neuropathology Sub-Core contributed to the tissue staining experiments), Grant C06RR029965 from the National Center for Research Resources, and by the National Institutes of Neurological Disorders and Stroke (NINDS) R01NS089664 and R01NS100874. JJW received supported from F31NS092264, TLS from F31NS095491, MA by a postdoctoral award from the National Ataxia Foundation (NAF), and AMB from F31NS101891. The content is solely the responsibility of the authors and does not necessarily represent the official views of the National Center for Research Resources or the National Institutes of Health.

